# Putting life history theory to the test - the estimation of reproductive values from field data

**DOI:** 10.1101/2022.03.09.483591

**Authors:** Mirjam J. Borger, Jan Komdeur, David S. Richardson, Franz J. Weissing

## Abstract

Quantifying fitness is important to understand adaptive evolution. Reproductive values are useful for making fitness comparisons involving different categories of individuals, like males and females. By definition, the reproductive value of a category is the expected *per capita* contribution of the members of that category to the gene pool of future generations. Life history theory reveals how reproductive values can be determined via the estimation of life-history parameters, but this requires intricate algebraic calculations. Recently, a more intuitive, pedigree-based method has become popular, which estimates the genetic contributions of individuals to future generations by tracking their descendants down the pedigree. Here we compare both methods. We implement various life-history scenarios (for which the “true” reproductive values can be calculated) in individual-based simulations, use the simulation data to estimate reproductive values with both methods, and compare the results with the true target values. We show that the pedigree-based estimation of reproductive values is either systematically biased (and hence inaccurate), or very imprecise. This holds even for simple life histories and under idealized conditions. In contrast, the traditional algebraic method estimates reproductive values with high accuracy and precision.

**Lay Summary:** To study evolution in empirical systems it is important to accurately and precisely measure fitness. Here, using simulations, we compare two methods for estimating fitness and test their accuracy and precision. One estimates life-history parameters and calculates a fitness proxy using complex algebra. The other method estimates genetic contribution to future generations by tracking descendants down the pedigree. We conclude that the pedigree method is unreliable, while the algebraic method is accurate and precise.

## Introduction

Quantifying fitness is important to understand how natural selection affects evolution. Estimating fitness in the wild is, however, a difficult task. Studies often measure survival, recruitment rate, or the number of offspring reaching independence as a proxy for fitness. In some well-monitored populations, lifetime reproductive success can be determined. Expected lifetime reproductive success is often a good proxy for fitness (Brommer et al., 2004), but even this measure is incomplete as it does not account for the rate of reproduction and the fact that different types of offspring (e.g. females and males; different size classes) cannot just be added up, as they have a different potential for spreading their genes to future generations.

Reproductive value (RV) (Fisher, 1930) is a comprehensive fitness measure that copes with these problems. The reproductive value of a certain class of individuals is defined as the expected contribution of an individual in that class to the future gene pool of the population (Grafen, 2006). Reproductive values are influenced by all kinds of life-history decisions, like age at first reproduction, trade-offs between survival and fecundity, risk-taking behaviour, or the kind of offspring produced (e.g. sons or daughters). RVs are a particularly useful tool in behavioural ecology, as they allow the evolutionary costs and benefits of fitness-relevant individual decisions to be quantified. For example, Tinbergen and Daan (1990) used an RV approach to predict the optimal clutch size in passerine birds on the basis of the trade-off between current and future reproduction. Reproductive values also play an important role in predicting the optimal sex ratio (Pen and Weissing, 2002) and in the evolutionary theories of senescence (Hamilton, 1966; Baudisch, 2005).

While the definition of reproductive value is straightforward, its measurement is not. As briefly reviewed below, RVs can be derived from a life-history model by solving an eigenvector equation that contains all relevant life-history parameters (Caswell, 1982). In line with this, most applications of RVs in empirical systems first estimate life-history parameters and subsequently obtain RVs by inserting these parameters in the eigenvector equation (Leverich and Levin, 1979; Schulman and Chapais, 1980; Tinbergen and Daan, 1990; Newton and Rothery, 1997; Pen et al., 1999; Bonduriansky and Brassil, 2002; van de Pol et al., 2007; Bouwhuis et al., 2012). Hereafter, we will call the RVs thus obtained “model-based reproductive values” (mRV). Obtaining a good estimate of reproductive values along these lines can, however, be difficult in natural systems, since it requires a good understanding of all relevant life-history transitions and because life-history parameters can often only be estimated with limited accuracy and precision.

Recently, reproductive values have been estimated with a more intuitive method, based on genetic pedigree data (Barton and Etheridge, 2011; Chen et al., 2019; Hunter et al., 2019; Reid et al., 2019). In this method, the average *per capita* number of descendants of the members of a certain class of individuals down the pedigree is used as an estimate of the reproductive value of that class. Hereafter, we will call the RVs thus obtained “pedigree-based reproductive values” (pRV). The pRV method closely reflects Fisher’s definition of reproductive value. This method requires the availability of sufficiently deep and complete pedigrees, but if such pedigrees are available it has the big advantage that no detailed knowledge of life-history transitions or estimates of life-history parameters are required. Therefore, the pRV method could potentially be a useful substitute for the mRV method. However, whether the pRV method matches the mRV method in accuracy and precision when applied to field data has yet to be determined.

In this theoretical study, we will compare the performance of both methods when applied to organisms with a relatively simple life history. To this end, we use individual-based simulations to produce several generations of individuals on the basis of a life-history model. The simulation data can be used to construct a pedigree (complete or incomplete) and to estimate life-history data such as survival probabilities and fecundities. Subsequently, reproductive values can be estimated by both methods (mRV and pRV) and compared with the “true” reproductive values (which are well-defined for the simulated populations). This allows us to judge the accuracy and precision of both estimates and to address questions like: how deep and complete must a pedigree be in order to get a reliable pRV estimate? How well must the life-history model be known, and how well do we need to know the life-history parameters in order to get reliable mRV estimates?

Before approaching the estimation problem, we first give some theoretical background by briefly reviewing some basal insights of life history theory.

## Theoretical considerations: why are reproductive values useful?

In a population with discrete, non-overlapping generations and only one type of individual (e.g. no sex differences), expected lifetime reproductive success (ELRS) is an adequate fitness measure, as alleles that enhance the lifetime reproductive output of their bearers have a selective advantage. The situation is different in populations with different classes of individuals (e.g. females and males; breeders and helpers; workers, soldiers and reproductives). In such cases, alleles inducing their bearers to produce a maximal number of offspring throughout their lifetime are not necessarily favoured. The reason is that different types of offspring may differ in the efficiency with which they spread genes to future generations. For example, in a population with a 4:1 female-biased sex ratio (4 females per male), the reproductive success per male is on average four times greater than the reproductive success per female (because each offspring has one mother and one father). Therefore, an allele inducing the lifetime production of two sons has a lower ELRS than an allele inducing the lifetime production of four daughters, but a higher “fitness”: it spreads more efficiently to future generations, as the expected reproductive success of two sons is twice the expected reproductive success of four daughters (for the given sex ratio). Similarly, adding up potential offspring does not result in a suitable fitness measure when generations are overlapping, and different age classes coexist. In this situation, the timing of reproduction matters: for example, it may be advantageous to produce offspring as early in life as possible (even if this leads to a lower lifetime production of offspring), as early-born offspring can more quickly contribute to the spread of their parental genes.

Quantifying fitness in a life-history context (i.e., in a population with different categories of individuals) is therefore not an easy task. Fortunately, life-history theory is a well-developed branch of evolutionary biology (Roff, 1992; Stearns, 1992; Caswell, 2001). A typical life-history model starts with a life-cycle graph (see Figure 1 for examples) that includes all life history “stages” (e.g. all age classes) and the transitions between these stages (e.g. survival to the next age class; production of offspring of age one). If the number of stages is finite, this graph can be translated into a “stage transition matrix” which in the case of three stages looks like this:

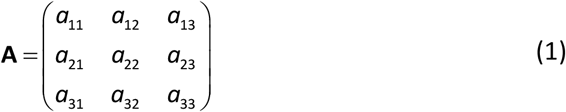

**Figure 1.**
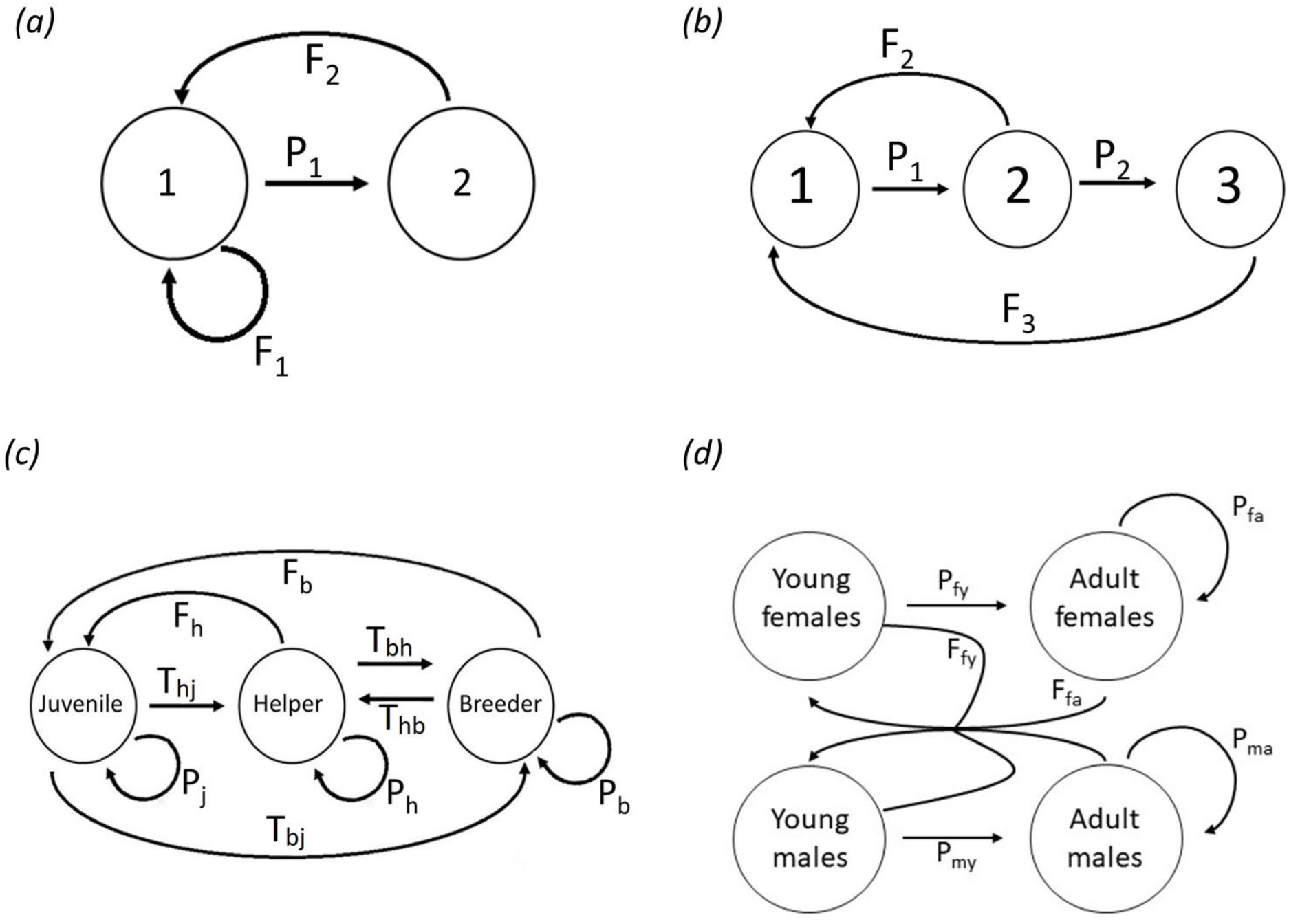
The four life-history scenarios considered in this study. **(a)** A two-stage life cycle with two age classes. One-year-old individuals survive to their second year with probability P_1_, second-year-old individuals die after reproduction. F_1_ and F_2_ denote the expected number of offspring (surviving to their first year) produced by one- and two-year-old individuals, respectively. **(b)** A three-stage life cycle with three age classes. Only two- and three-year-old individuals can reproduce (with expected fecundity F_2_ and F_3_), and one- and two-year-old individuals survive to the next year with probabilities P_1_ and P_2_, respectively. **(c)** A three-stage life cycle with behavioural stages. A juvenile can either stay juvenile for another time unit (probability P_j_), promote to helper status (probability T_hj_), promote to breeder status (probability T_bj_), or die. A helper can either stay a helper for another time unit (probability P_h_), promote to breeder status (probability T_bh_), or die. A breeder can stay a breeder (probability P_b_) or die. Both helpers and breeders can reproduce with expected fecundity (number of offspring surviving to the juvenile stage) F_h_ and F_b_, respectively. **(d)** A four-stage life cycle including differences between sexes. Young females and young males survive to become an adult with probabilities P_fy_ and P_my_ respectively. Adult females and adult males survive to the next year (and stay in the same state) with probabilities Pfa and Pma respectively. Young females and adult females have expected fecundities of Ffy and Ffa. They randomly pick a mate from the population of males. A newly produced offspring becomes a male with probability *s* and a female with probability 1-*s*, where *s* is the primary sex ratio.

Matrix element *a_ij_* is the *per capita* contribution of a member of stage *j* to the number of individuals in stage *i* in the next time step. For example, the stage transition matrix of an age-structured population with three age classes has the special form of a “Leslie matrix”:

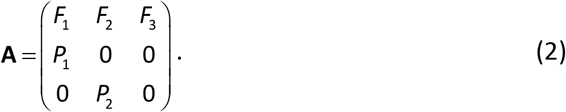

Here, *P*_1_ is the probability to survive from age class 1 to age class 2, *P*_2_ is the probability to survive from age 2 to age 3, and *F*_1_, *F*_2_, and *F*_3_ is the expected number of surviving offspring (surviving till age class 1) produced per individual of the respective age class.

Life-history matrices are a very useful tool, as they determine the dynamics of a stage-structured population (Stearns, 1992). In most cases, such a population will converge to a “stable stage distribution” (in the case of an age-structured population: a stable age distribution; in the case of a population with two sexes: a stable sex ratio), that is, to a state where the ratio between the number of individuals in the stages (*n*_1_: *n*_2_: *n*_3_) does not change anymore. Once the stable stage distribution is reached, the population as a whole grows with a characteristic “population growth factor *λ*”. When *λ*=1, the population size will stay constant over time, while the population size will increase exponentially when *λ*>1 and decrease exponentially when *λ*<1.

The population growth factor *λ* is the most fundamental fitness measure in a stage-structured population (Brommer, 2000): if we consider a population with transition matrix **A** and corresponding *λ*, a mutant inducing a slightly different transition matrix **A_m_** with growth factor *λ_m_* can be expected to successfully invade the population if *λ_m_* > *λ*, while mutants with *λ_m_* < *λ* will go extinct. Mathematically, *λ* is the “dominant eigenvalue” of the stage-transition matrix **A**, and there are recipes to calculate *λ* for a given matrix **A**. Unfortunately, the eigenvalue equation determining *λ* is an “implicit” equation (in case of an age-structured population the so-called “Euler-Lotka equation” (Otto and Day, 2007)), which even for simple life histories can only be solved numerically and does not provide much insight into the factors governing evolution. It is therefore useful to look out for an alternative fitness measure.

At this point reproductive values come into play. Conceptually, the “reproductive value” *v_i_* of a stage *i* is the expected genetic contribution of a member of stage *i* to the gene pool of future generations (Grafen, 2006). Mathematically, the vector **V**^T^ = (*V*_1_,*V*_2_,*V*_3_) of reproductive values satisfies the equation **v**^*T*^ · **A** = *λ*·**v**^*T*^, which means that it is a “left eigenvector” with respect to the dominant eigenvalue *λ*. Reproductive values are useful for making evolutionary predictions in two different ways. First, in many life-history contexts, including those of age-structured populations, natural selection tends to maximize the reproductive value of each age class (Schaffer, 1974). This is the basis of dynamic programming (Houston and McNamara, 1999), a powerful technique for solving difficult problems in behavioural ecology, such as finding the optimal allocation of resources to various activities, like growth and reproduction. Second, and more important for us, the reproductive values in a given “resident” population can be used to calculate the “selection gradient” *∂λ*/*∂x*, which tells us whether the population growth factor *λ* (the most fundamental measure of fitness) will increase or decrease with a change in a (behavioural) strategy *x*. In other words, the selection gradient tells us the direction of selection: if *∂λ*/*∂x* is evaluated at the strategy *x*^*^ of the resident population, a positive selection gradient indicates that larger values *x* > *x*^*^ are selectively favoured, while smaller values *x* < *x*^*^ are favoured in the case of a negative gradient. Although *λ* is very difficult to calculate as a function of *x*, the sign of the selection gradient can relatively easily be determined by the following equation (Taylor, 1990; Taylor and Frank, 1996; Brommer, 2000; Pen and Weissing, 2000b, 2002; Otto and Day, 2007; Lion, 2018):

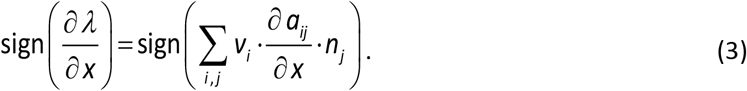

Here, the *v_i_* and the n_*i*_ are the reproductive values and the relative abundances of the various stages in the resident population (at demographic equilibrium) and the derivatives are evaluated at *x* = *x*^*^.

Why is an equation like this useful? Assume that we want to know whether, in an age-structured population, selection will favour an increase in the reproductive effort at age *i*. To this end, let *x* denote a behavioural strategy that enhances the reproductive output at age *i* with a rate *∂F_i_*/*∂x* = *b* but reduces survival to the next age with a rate *∂P_i_*/*∂x* = −*c*, where *b* and *c* indicate the “benefit” and the “cost” of an increase in *x*. Now the sum on the right-hand side of (3) has only two terms, and the sign of the selection gradient is readily obtained

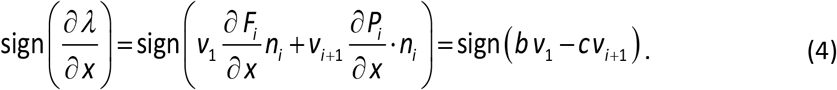

Hence, the selection gradient is positive if *bv*_1_ > *cv*_*i*+1_, or equivalently:

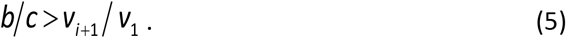

This inequality is known as the “asset protection principle”: the age classes with the highest future fitness expectation *V*_*i*+1_ (i.e. those age classes that have “much to lose”) should be least inclined to take the survival risks associated with increasing the reproductive effort (see Wolf et al., 2007 for various implications of this principle).

As illustrated by these calculations, reproductive values are useful tools in translating intricate fitness considerations into transparent cost-benefit comparisons (see Pen & Weissing 2000a, 2000b, 2002 for other examples). Cost-benefit considerations are further simplified by the fact that reproductive values are *relative* attributes that are only determined up to a constant factor. This implies that they can be normalized in the most convenient manner, for example by equating one of the reproductive values to one (e.g., *V*_1_ = 1, which would simplify eqn (5)) or by equating the sum of all reproductive values to one. Finally, we would like to remark that classical life history theory is ecologically not consistent, in that its models predict exponential population growth (when *λ* > 1) or exponentially population decline (when *λ* < 1). To cope with this problem, density dependence needs to be incorporated into a life history model. This has to be done in an explicit manner, as the form of density dependence can have major implications for the course and outcome of evolution (Mylius and Diekman, 1995; Pen and Weissing, 2000b). Below we illustrate how to introduce density dependence in a life-history model. Rather than complicating the model, this actually simplifies calculations, as the inclusion of density dependence has two benefits: (a) it allows the assumption that *λ* = 1 at ecological equilibrium and (b) it reduces the dimensionality of the parameter space (thus making it easier to classify the evolutionary outcomes).

## Methods

### Simulations for the pedigree-based estimation of reproductive values

All individual-based simulations were based on a specific life-history scenario. Each scenario is specified by a set of parameters (such as the age-dependent survival probabilities *P_i_* and fecundities *F_i_* characterizing an age-structured population). For a given state *X*, all individuals in that state have the same state-specific parameters. Accordingly, all individuals in that state have the same (expected) state-specific reproductive value. Time proceeds in discrete steps, where one time step corresponds to a reproductive season. During a time step *t*, each individual is in a certain state (either an age class or a breeding state), which can change from one time step to the next. Reproduction takes place at the start of each time step. If, according to the life history scenario, *F_x_* is the expected fecundity in state X, the individuals in that stage produce *F_x_* offspring on average, while the actual number of offspring produced per individual is drawn from a Poisson distribution with mean *F_x_*. Offspring enter the population at time *t*+1, and they belong to age class 1 in the age-based scenarios and to the juvenile state in the breeder-helper scenario. After reproduction, all individuals present at time *t* change their state stochastically (including death and staying in the same state), according to the probability assigned to their state by the life-history model. Each simulation was repeated for a fixed number of time steps *T* (typically *T*=100, of which the first 20 time steps are shown in the figures). For each life-history scenario, simulations were run for various parameter combinations, and in each case the simulations were repeated at least 100 times. To ensure that our results were not biased by start-up effects, we initialized all populations in demographic equilibrium. Our results are therefore not examples of “transient dynamics” (Hastings, 2004; Hastings et al., 2018). This was confirmed by additional simulations that were first run for thousands of time steps (thus ensuring equilibration) before the measurements were started.

We used a “gene-dropping” approach (MacCluer et al., 1986) to estimate pRVs. At the start of the simulation (*t*=0), every individual is endowed with a unique marker (corresponding to a unique allele at a locus with infinite alleles). To ensure that gene dropping happened in demographic equilibrium, we first let each simulation run for 100 time steps (from t=-100 until t=-1) before gene dropping was initiated (at t=0). At each reproduction event, the offspring inherit their marker from their parent. This allowed us to identify the descendants of each individual of the initial population in all future time steps. For each life history state *X* and each time *t*, we determined the average *per capita* number of descendants at time *t* for those individuals in the initial population that started in state *X*. We interpret this number, *pRV_X_*(*t*), as the pedigree-based estimate of the RV of individuals in state *X*: according to the definition of RV (“*per capita* contribution to the gene pool of future generations”) *pRV_X_*(*t*) should, for large *t*, approximate the “true” reproductive value of state *X*. We then normalize all reproductive values so that the RV of the “first” state (age class 1 in the age-structured scenarios; juvenile state in the helper-breeder scenario; young female in the sex-and-age-structured scenario) is scaled to 1. To achieve this, the values *pRV_X_*(*t*) were divided by *pRV*_1_(*t*), the estimated reproductive value of the first state. For simplicity, we assumed that marker inheritance is based on asexual reproduction in most simulations, and that individuals are haploid. However, in the simulations where sex was included, sexual reproduction and diploidy was assumed. As shown in Appendix A in the Supplement, the expected value of *pRV_X_*(*t*) can be calculated mathematically.

To prevent exponential growth or exponential decline of the simulated populations, we made the reasonable assumption that population sizes are kept in check by density dependent processes. Therefore, we added density regulation by assuming that fecundity is density dependent:

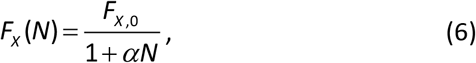

Where *F*_*X*,0_ is the baseline fecundity in state *X*, *N* is population size and α indicates the intensity of density dependence. In other simulations (not shown), we also implemented other forms of density regulation (via density-dependent fecundity of only a single state or via density-dependent survival of one or all states); in all cases, the results agreed with those shown below. Unless indicated otherwise, we chose the parameter α in such a way that the population size of the resulting stationary population was N=1,000. Note that this means that about 1,000 individuals are present at every time step. If *T* is the depth of the pedigree and *E* is the life expectancy of individuals, this implies that the pedigree encompasses about (*T/E*) ·1,000 individuals. Thus, in all our simulations with a time horizon *T* = 20 at least 10,000 individuals were included in the simulation-based pedigree.

For the model-based estimation of reproductive values (mRV), the life-history parameters had to be estimated from the life-history events (survival, death, state transition, offspring production) observed in the simulations. To estimate *mRV_X_*(*t*) for a certain time *t*, we used all events observed on individuals in state *X* up to time *t* to estimate the life-history parameters relevant for that state by averaging. For example, the fecundity *F_x_* in state *X* was estimated by the average number of offspring produced in state *X* (per time step) by all individuals that had ever been in state *X* until time *t*. Subsequently, the *mRV_x_*(*t*) were estimated by calculating the left eigenvector of the matrix **A**(*t*) that is characterized by all parameter estimates up to time t.

### Four scenarios

In this manuscript we focus on four life history scenarios, which will be described in detail below. The life-cycle graphs of these scenarios can be found in figure 1 and the corresponding transition matrices and calculations on reproductive value can be found in Appendix B.

#### Scenario 1 A population with two age classes

This simple scenario, with only two life-history states and three life-history parameters, is characterized by Figure 1(a). Both one- and two-year-old individuals can reproduce (with fecundity F_1_ and F_2_). One-year-old individuals survive to their second year with probability P_1_; all individuals die after their second year. A simple calculation (see Appendix B) reveals that a stationary population (*λ* = 1) is only achieved if the life-history parameters satisfy the requirement:

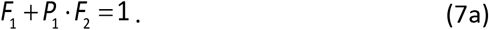

This equation makes intuitive sense: the left-hand side corresponds to the expected lifetime production of surviving offspring, which needs to be 1 in a stationary population. Condition (7a) for “ecological consistency” will not be satisfied for arbitrary values of the life-history parameters. However, it will eventually be satisfied, as, according to eqn (6), the fecundities decline with population density.

As shown in Appendix B, the “true” reproductive values are given by:

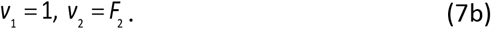

#### Scenario 2 A population with three age classes

This scenario, with three life-history states and four life-history parameters, is described in Figure 1(b). Two- and three-year-old individuals can reproduce (with fecundity *F*_2_ and *F*_3_). One- and two-year-old individuals can survive to their next year (with probabilities *P*_1_ and *P*_2_); all individuals die after their third year. Now the condition for a stationary population and the reproductive values in that population are given by:

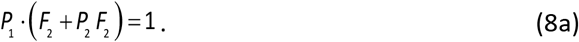

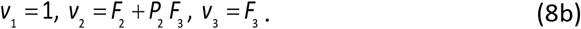

Ecological consistency (a stationary population) is ensured by eqn (6).

#### Scenario 3 A population with three behavioural classes

This scenario, which is characterized by Figure 1(c), includes three life-history states (juveniles, helpers, breeders) and a full set of nine life-history parameters. All newly produced individuals start as juveniles. A juvenile either stays in juvenile state for another time period (with probability *P_j_*), or it becomes a helper (with probability *T_hj_*) or a breeder (with probability *T_bj_*), or it dies (with probability 1−*P_j_*−*T_hj_*−*T_bj_*). Likewise, a helper either stays in the helper state for another period (probability *P_h_*), or it becomes a breeder (probability *T_bh_*), or it dies. A breeder either stays in the breeding state for another time step (probability *P_b_*), or it loses its position and becomes a helper (probability Thb.), or it dies. Helpers and breeders can both produce offspring (fecundities *F_h_* and *F_b_* per time step).

#### Scenario 4 A population with both sex and age classes

This scenario, which is characterized by the life-cycle graph in Figure 1(d), contains four life-history states: young females, young males, adult females and adult males. Newly produced individuals start as young females or young males. The sex of an offspring is assigned at random, where the probability *s* of becoming a male corresponds to the primary sex ratio. The parameter *s* is the same for all individuals and constant throughout time. Young females survive to become adults with probability Pfy, and young males survive to become adults with probability Pmy. Adult females and males survive to the next time step with probabilities Pfa and Pma respectively. Both young and adult individuals can reproduce. We assume that young and adult females produce on average *F*_fy_ and *F*_fa_ offspring per time step, respectively, where these fecundities are density dependent according to (6). For simplicity, we assume that each male has the same probability of siring any given offspring: whenever an offspring was produced, a father was drawn at random from the set of all (young and adult) males.

### Technical note

Simulations were ran in Visual Studio Enterprise 2019 (version 16.8). Figures were made using R 3.4.1 (R Core Team, 2021), with the packages ggplot2 (Wickham, 2016) and cowplot (Wilke, 2019).

## Results

### Stochasticity in the number of descendants

For simulations based on Scenario 2 (three age classes), Figure 2 illustrates different sources of stochasticity in the number of descendants per individual. Figure 2a shows that, within a single simulation, stochasticity in individual reproductive success is extensive. For example, two of the three individuals that started in age class 1 have not left any descendants after 20 years, while the third individual (Individual 2) leaves about 20 descendants. Similar patterns are seen for the other age classes. Estimating RV based on a single individuals is therefore highly inaccurate, since these values mainly reflect stochasticity, rather than the capability of a class of individuals to spread their genes to future generations. When using estimates based on the averages of entire cohorts (or behavioural groups in later scenarios), this stochasticity partially gets averaged out. Yet, as shown in Figure 2b, each simulation still creates a unique pattern of numbers of descendants per age group. To get an impression of this variation, we ran 100 replicate simulations, resulting in Figure 2c, which shows the median values and the 50% and 90% central values of these 100 simulations. Figure 2c shows that the theoretically expected values (dashed lines) are approached in a “zig-zag” manner. As shown in Appendix A, this is not caused by “transient dynamics” (short-term deviation from demographic equilibrium), but an intrinsic property of the gene-dropping dynamics. Although the expected values are approached on a longer-term perspective, the simulations still differ considerably in the number of descendants per cohort.

**Figure 2.**
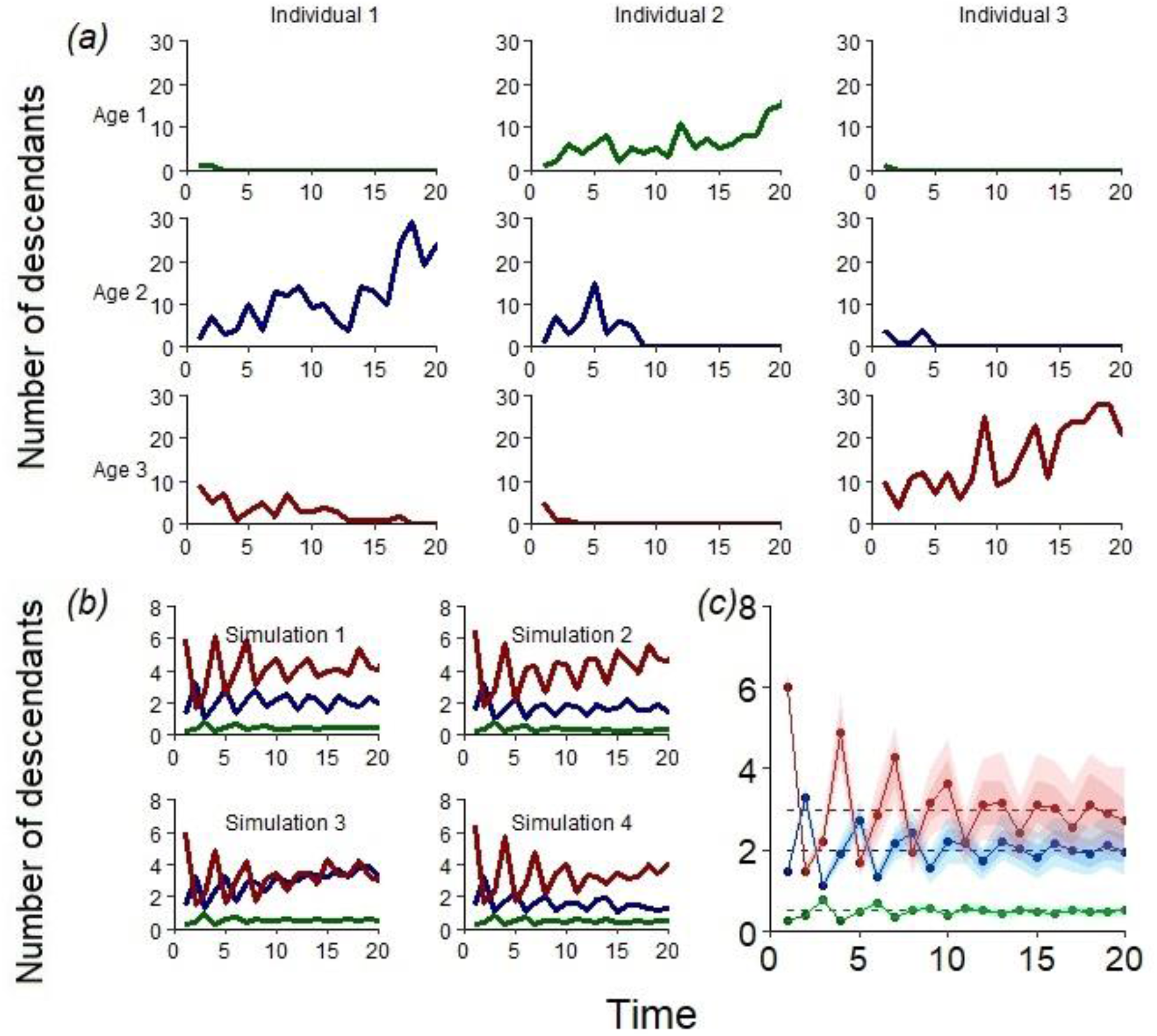
Stochasticity in the number of descendants. Example simulations based on Scenario 2 (three age classes; Fig. 1b) illustrate stochasticity within and across simulations. **(a)** Number of descendants at a given time step (0 to 20) from nine individuals (three per age class) that were present in the same simulation at time zero. **(b)** For four replicate simulations, the panels show the number of descendants at a given time step of the three cohorts at the start of the simulation (t=0): one-year olds (green), two-year olds (blue), and three-year olds (red). **(c)** Summary of 100 simulations. Dashed horizontal lines correspond to the number of descendants expected by the “true” RVs. The dots connected by solid lines indicate the median of the 100 simulations. The 50% (respectively 90%) central values of the simulations are represented by the darker (respectively lighter) shaded area around the medians. Parameter values: P_1_=0.25, P_2_=0.5, F_2_=1 and F_3_=6.

### RV estimates in Scenario 2

In Figure 3, we show the RV estimates of Scenario 2. From now on, we follow the standard practice in life history theory and report normalized reproductive values, where the reproductive value of stage 1 is set to one (i.e., *V*_1_=1). In Scenario 2, we can therefore focus on age classes 2 and 3. In Figure 3a-c it can be seen than individual simulations (comparable to estimating RV in a field population) show considerable variation in pRV estimates, with some estimates being highly inaccurate. In contrast, the mRV estimates are both accurate (they closely approximate the true RVs) and precise (they do not show much variation between replicate simulations). The summary of 100 replicate simulations in Figure 3d-f shows that these conclusions are hold in general: for Scenario 2, mRVs provide an accurate and precise estimate of true RVs, while pRVs are quite inaccurate (in particular in the first 6 time steps) and/or unprecise (in particular in the later time steps. Note that one replicate simulation corresponds to one field population of size N = 1,000, of which a complete pedigree is available. Figure 3 therefore exemplifies that, at least in Scenario 2, pedigree-based estimation of RVs is problematic: On a short time horizon, the pRV estimates agree with each other reasonably well, but they deviate substantially and systematically from their target value, the “true” RVs. These deviations occur despite of the fact that the populations are reasonably large, and all simulations take place at demographic and ecological equilibrium. In Appendix A of the Supplement, we show mathematically that, for small values of *t*, large deviations are always to be expected. On a longer time horizon (t>10) the median pRVs approximate the true RVs, but now the pRVs differ considerably between simulations.

**Figure 3.**
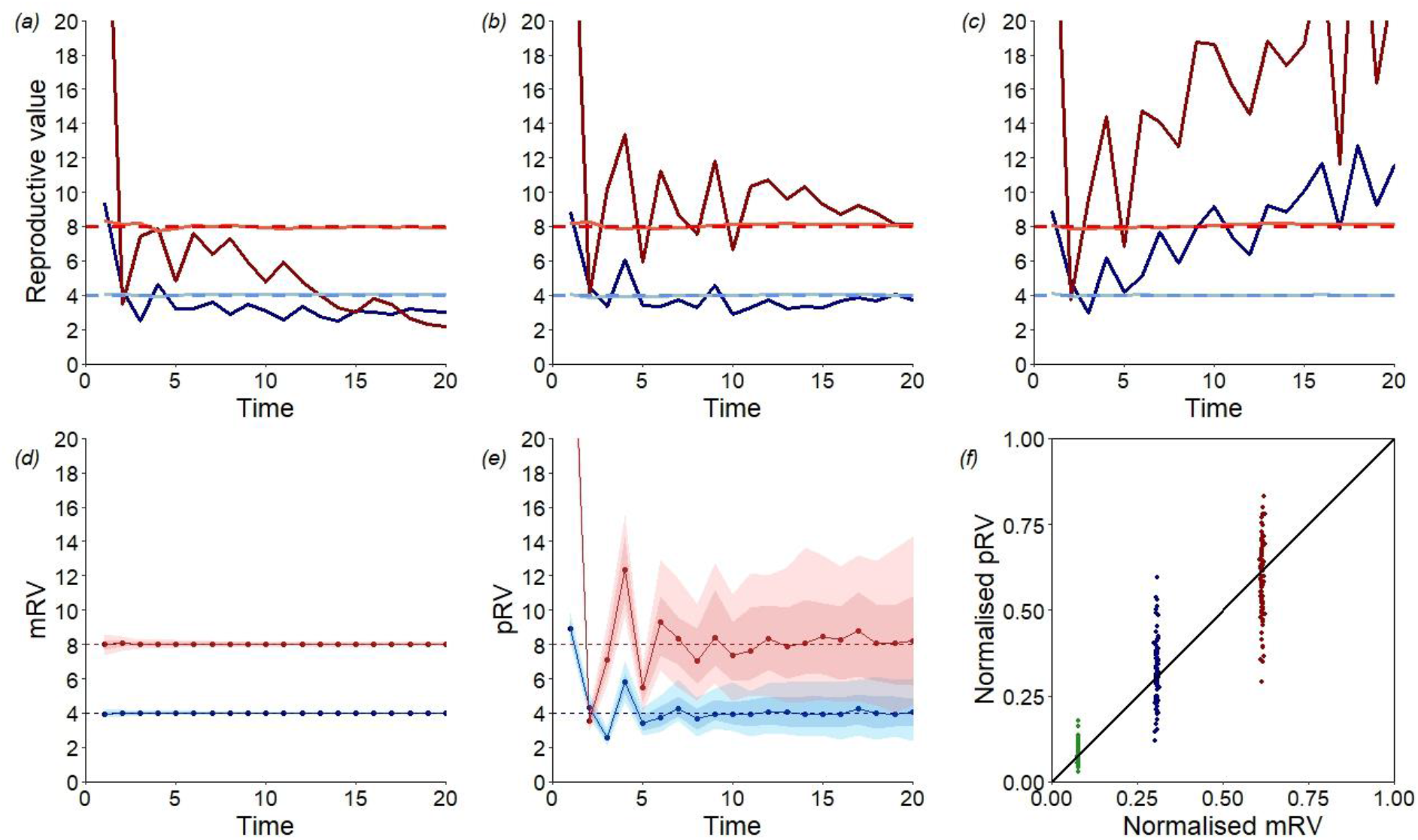
mRV and pRV estimates in Scenario 2 (3 age classes) **(a)-(c)** Three replicate simulations (comparable to field populations) illustrating the diversity of outcomes. The dark-coloured lines (dark blue for two-year-olds and dark red for three-year-olds) represent pRV estimates, while the light-coloured lines (light blue and bright red for two- and three-year-olds, respectively) represent mRV estimates. The dashed lines (blue for two-year-olds and red for three-year-olds) represent the “true” values of the RVs. The simulations exemplify that (a) pRVs can substantially underestimate true RVs; (b) pRVs can approach true RVs; and (c) pRVs can substantially overestimate true RVs. In contrast, mRVs approximate true RVs very well in all three cases. **(d,e)** Summary of mRV and pRV estimates from 100 replicate simulations. In both cases, the dashed horizontal lines correspond to the true RVs. The dots connected by solid lines indicate the median of the 100 simulations. The 50% (respectively 90%) central values of the simulations are represented by the darker (respectively lighter) shaded area around the medians. **(f)** The pRV estimates of 100 simulations are plotted against the accompanying mRV estimates, all from time step t=20. To show the relationship between the pRV and the mRV more clearly, we normalized the RVs in such a way that their sum equates to 1 (instead of normalizing *V*_1_ to 1). Here, the green dots represent the RV estimates of one-year-olds, the blue dots represent RV estimates of two-year-olds and the red dots represent RV estimates of three-year-olds. Parameter values: P_1_=0.25, P_2_=0.25, F_2_=2 and F_3_=8.

### RV estimates in Scenario 1

Figure 4 shows that even for the extremely simple life history of Scenario 1, pRV is either a systematically biased (on a short time horizon) or imprecise (on a longer time horizon) estimator of RV. In contrast, “true” RV is, from the start, estimated with accuracy and precision by mRV.

**Figure 4.**
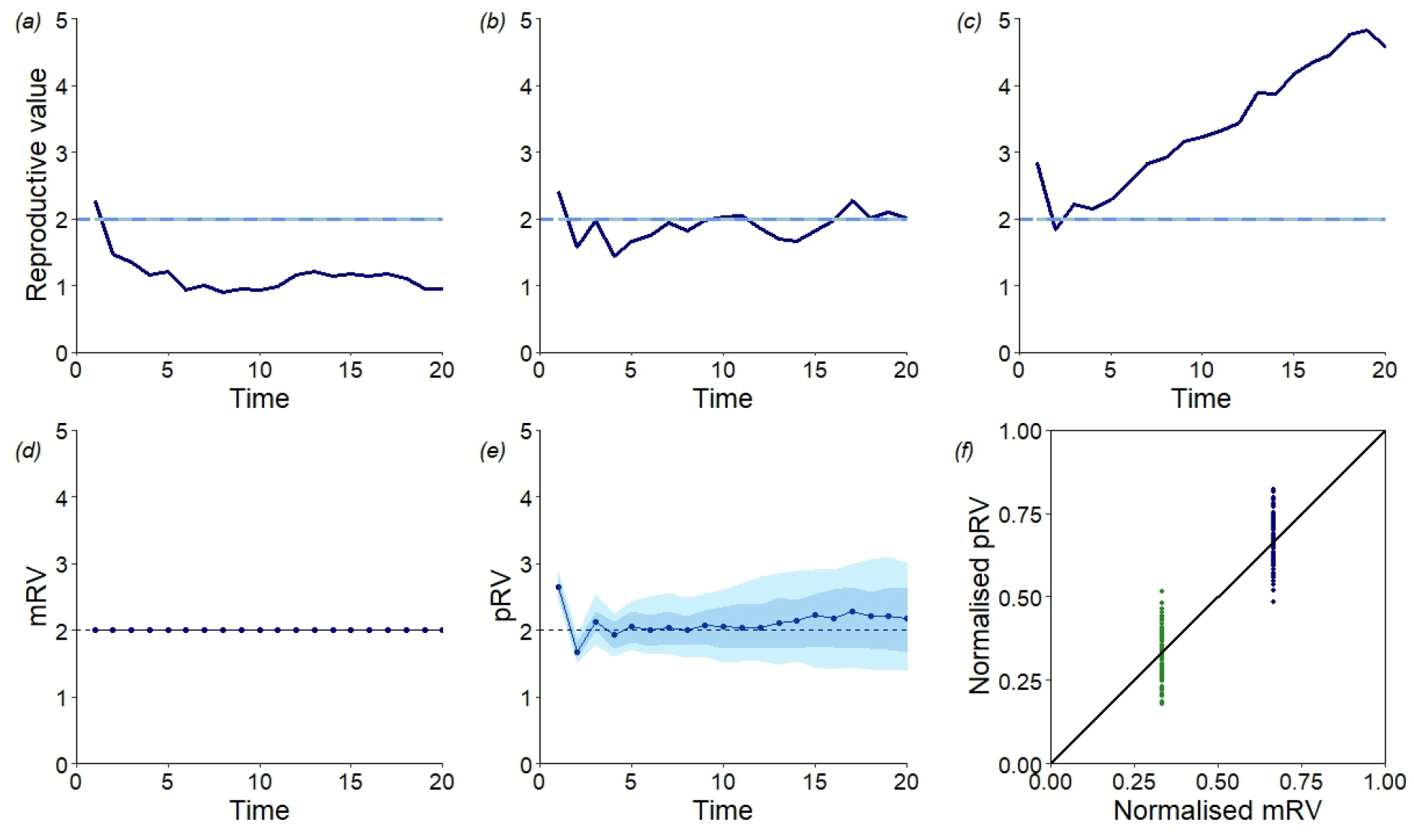
mRV and pRV estimates in Scenario 1 (2 age classes). **(a)-(c)** Three example simulations (comparable to field populations). The dark blue lines represent pRV estimates (for two-year-olds), while the light blue lines represent mRV estimates. The dashed lines represent the “true” values of the RVs. The simulations exemplify that (a) pRVs can substantially underestimate true RVs; (b) pRVs can approach true RVs; and (c) pRVs can substantially overestimate true RVs. In contrast, mRVs approximate true RVs very well in all three cases. **(d,e)** Summaries of mRV and pRV estimates from 100 replicate simulations. In both cases, the dashed horizontal lines correspond to the true RVs. The dots connected by solid lines indicate the median of the 100 simulations. The 50% (respectively 90%) central values of the simulations are represented by the darker (respectively lighter) shaded area around the medians. **(f)**The pRV estimates of 100 simulations are plotted against the accompanying mRV estimates, all from time step t=20. To show the relationship between the pRV and the mRV more clearly, we normalized the RVs in such a way that their sum equates to 1 (instead of normalizing *V*_1_ to 1). Here, the green dots represent the RV estimates of one-year-olds and the blue dots represent RV estimates of two-year-olds. Parameter values: F_1_ = 0.5, P_1_ = 0.25, F_2_ = 2.

### RV estimates in Scenarios 3 and 4

Figure 5 shows that the problems with pRV estimates also arise in other life-history scenarios. In Scenario 3 (Fig. 5a), the pRV estimates can be two or even three times as large as the true RVs in a considerable percentage of the simulations. Although the initial “zig-zag pattern” observed in Scenarios 2 and 3 does not appear in Scenario 4 (Fig. 5b), pRV remains an inaccurate estimator of true RV in the initial period; in fact, the systematic bias in this estimate only disappears after a long period (in this case 10 time steps). Hence, the simulations in Fig. 5 confirm our earlier conclusions: in an initial time period, the pRV values (including their median) differ substantially and systematically from the true RVs. On a longer time horizon, the accuracy of pRV increases (the median pRVs approach the true RVs), but now the estimates are very imprecise, as the individual simulations differ considerable from each other. As one simulation corresponds to a single field study, this results in the unfortunate conclusion that pedigree-based estimation of RVs is either inaccurate (in case of a shallow pedigrees including relatively few time steps) or imprecise (in case of deep pedigrees), even if the study population is relatively large (*N*= 1,000 in the simulations shown thus far), well-mixed, and in demographic and ecological equilibrium.

**Figure 5.**
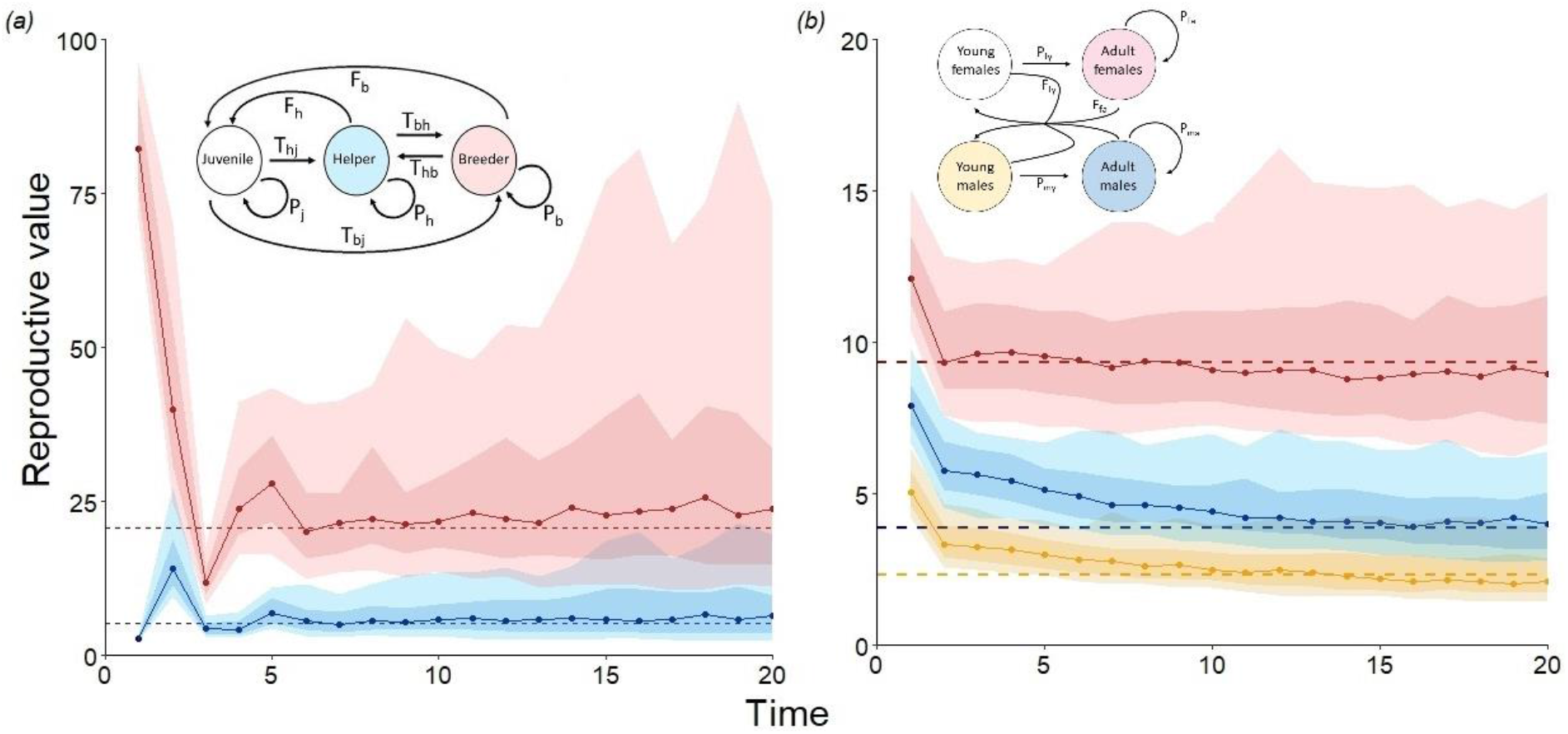
Accuracy and precision of pedigree-based estimates of RV in two more complex scenarios. Pedigree-based estimation of two (normalised) RVs in 100 replicated simulations based on **(a)** Scenario 3 (juvenile-helper-breeder; Fig. 1(c)), and **(b)** Scenario 4 (young and old males and females; Fig 1(d)). The inserts show the life-cycle graphs. The “true” RVs are represented by dotted lines. Dots connected by solid lines indicate the median pRV of the 100 simulations; the 50% (resp. 90%) central values of the 100 pRVs are represented by the darker (respectively lighter) shaded area around the medians. In (a) blue represent the RV values of helpers and red represent the RV values of breeders. In (b) yellow represent the RV values of young males, while red and blue represent the RV values of adult females and males, respectively. Please note the difference in scale on the y-axis. Parameter values (a): F_j_=0, F_h_=0.05, F_b_=12.5, T_hj_=0.15, P_h_=0.2, T_hb_=0, T_bj_=0.01, T_bh_=0.2, P_b_=0.4 and (b): P_fy_=0.1, P_fa_=0.85, P_my_=0.5, P_ma_=0.9, F_fy_=0.095, F_fa_=2, s=0.3.

### Effects of population size and time scale

In Appendix C of the Supplement, we also show the effects of population size and a longer time horizon on pRV estimates. When population sizes are small, pRV estimates are strongly affected by demographic stochasticity. After a while, all individuals present at a later time step are descendants of just one individual of the initial population. Accordingly, in each simulation, eventually pRV is equal to zero for all but one of the categories considered. Therefore, in small populations, the median pRV of 100 simulations converges to zero relatively rapidly (Supplementary Figure C1(a)). Furthermore, the variation of the pRV estimates across simulations increases with the time horizon (Supplementary Figs C1 and C2). In other words, the precision of pRV decreases with the depth of the pedigree.

### Evolutionary predictions based on RV

Above, we have seen that reproductive values play an important role in predicting the course and outcome of life history evolution. One might speculate that the large variance of pRV values that we observed in replicate simulations is associated with a corresponding variance in evolutionary outcomes. To check this, we added an evolving parameter *x* (corresponding to reproductive effort) to our Scenario 1 model with two age classes. To this end, we added a gene locus to the model, with a continuum of alleles, ranging from 0 to 1. Individuals with allele *x* at this locus have life-history parameters 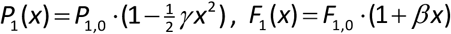, and *F*_2,0_(*x*) = constant, where β and γ are positive. Hence, one-year olds with a larger reproductive effort *x* have a higher reproductive output *F*_1_, but a lower survival probability *P*_1_. We assume that *x* is transmitted from parent to offspring, subject to rare mutations with small effect size (mutation rate 0.01, mutational variance 0.0025; see Netz et al. 2022). In the course of the generations, *x* should evolve to a value that optimizes the balance between current and future reproduction. With the help of reproductive values, the optimal reproductive effort can easily be calculated. At evolutionary equilibrium, the selection gradient is zero, which according to eqn (4) corresponds to the equation 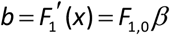 and 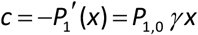, while *v*_2_/*v*_1_=*F*_2,0_ in view of eqn (7b). Inserting all these terms and solving for *x*, we obtain the optimal reproductive effort

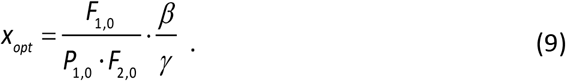

Figure 6 demonstrates that, for a set of parameters yielding the optimal value *x_opt_*=0.5, individual-based simulations do indeed converge to this value, and that they stay close to this value, irrespective of whether the pedigree-based estimate *pRV*_2_/*pRV*_1_ of *v*_2_/*v*_1_ is much larger or much smaller than the “true” value *F*_2,0_. We conclude that (a) reproductive values are indeed useful for making evolutionary predictions; but that (b) pedigree-based estimates are too imprecise to be reliable predictors of the evolutionary outcome.

**Figure 6.**
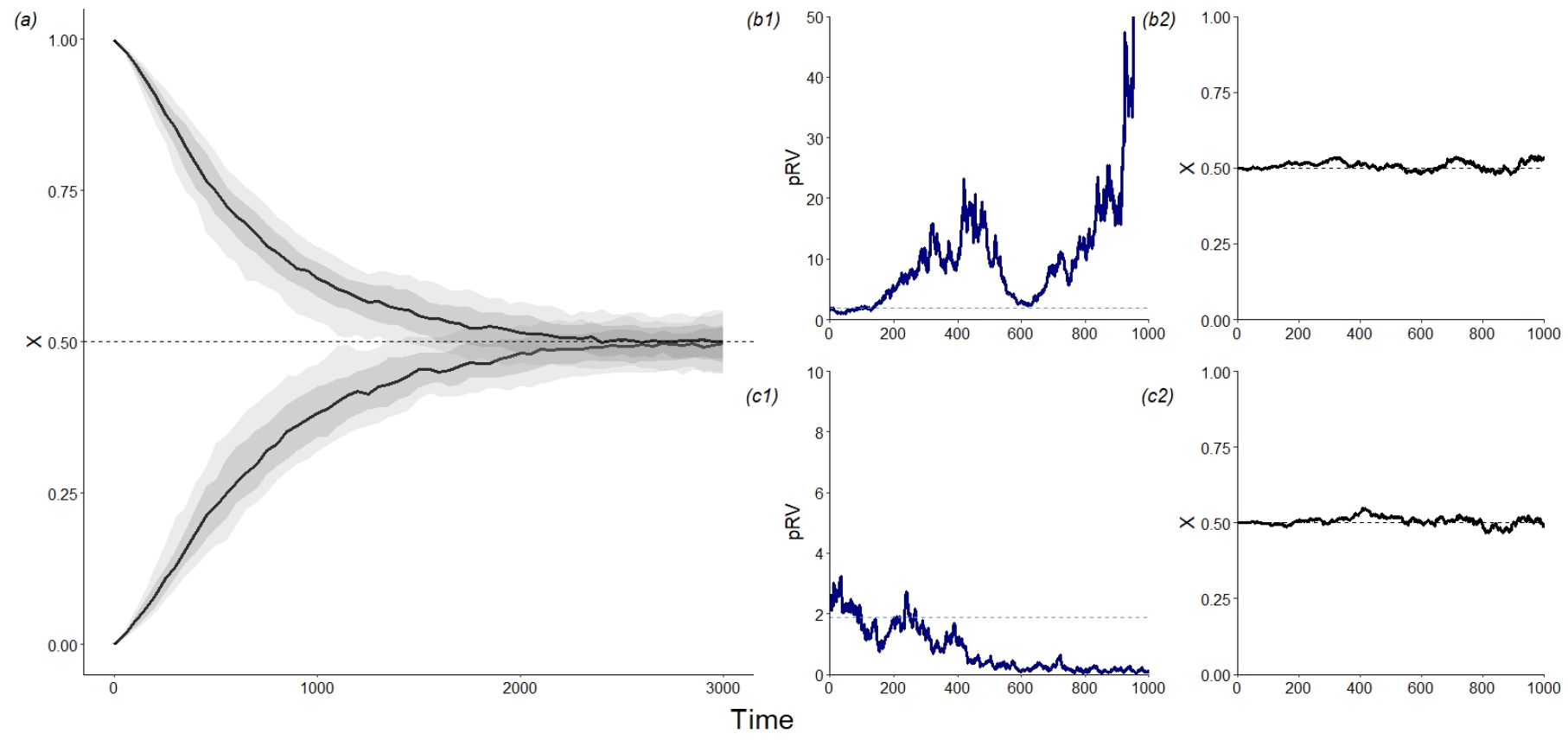
Evolutionary predictions based on reproductive values. **(a)** Evolution of reproductive effort *x* in Scenario 1 (two age classes) in two sets of 100 individual-based simulations; one set starting at *x* = 0 and the other starting at *x* = 1. shows the trajectory of x over time for 200 simulations (100 that started with a monomorphic population where *x* = 0, and 100 that started with a monomorphic population where x=1). The solid lines indicate the median *x*-values of these simulations and the 50% (resp. 90%) central values of x are represented by the darker (resp. lighter) shaded areas around the medians. The dashed line indicates the optimal reproductive effort, which, based on the “true” reproductive values, is equal to *x_opt_*=0.5, according to eqn (9) and the parameter setting *P*_1,0_ = 0.25, *F*_1,0_ = 0.5, *F*_2,0_ = 2, *β* = 0.5, *γ* = 1. **(b)** and **(c)** show two example simulations that both start at equilibrium (*x* = 0.5). The evolutionary trajectories of *x* ((b2) and (c2)) stay close to this equilibrium value, while the pRV ratio *pRV*_2_/*pRV*_1_ either overestimated (b1) or underestimated (c1) the “true” RV ratio *v*_2_/*v*_1_ (dashed line).

## Discussion

Reproductive values are a very useful theoretical tool, as they can help to answer questions like: what is the optimal clutch size (Tinbergen and Daan, 1990)? What is the optimal investment in male versus female offspring (Fisher, 1930; Pen and Weissing, 2002)? Under which circumstances should offspring stay on their natal territory and help their parents raise additional offspring (Pen and Weissing, 2000c)? However, we need to be able estimate RVs with accuracy and precision if we are to use them to test predictions of life-history theory.

Reproductive values are increasingly estimated by making use of pedigree information from long term studies (e.g. Barton and Etheridge, 2011; Chen et al., 2019; Hunter et al., 2019; Reid et al. 2019). In essence, the *per capita* number of descendants of the members of a certain life-history stage is used to estimate the reproductive value of that stage. This method is intuitively appealing, as it closely reflects Fisher’s (1930) definition of “reproductive value”. However, our study clearly reveals that the method has important drawbacks in practice, even if the pedigree includes many individuals (hundreds or even thousands) per time step. The analytical arguments (Appendix A) and simulations outlined in the results show that pedigree-based estimates are strongly and systematically biased (and therefore inaccurate) if the pedigree encompasses a short time horizon (say, *t* < 10). On a longer time horizon, the median pRVs of 100 replicate simulations match the “true” RVs reasonably well, but the individual simulations tend to diverge considerably from each other with increasing “depth”of the pedigree. As one simulation corresponds to one field study, this implies that the pedigree-based estimate of RV is typically way “off target” if it is only based on one, or a small number of, field studies.

Our findings concur with the patterns reported in natural populations. In both Soay sheep (*Ovis aries*, Hunter et al., 2019) and Florida Scrub-Jays (*Aphelocoma coerulescens*, Chen et al., 2019) estimates of RV based on individuals did not stabilise, just as in our simulations. Moreover, the pedigree-based estimates showed a similar initial “zig-zag” pattern as in our study (e.g. Fig. 2b, c, 3e, 4e, 5a). Last, but not least, when the estimation was repeated for the same population by applying the gene-dropping method to different cohorts, the RV estimates obtained varied a lot (Hunter et al., 2019).

Our study confirms that the more traditional, model-based calculation of reproductive values on the basis of estimates of the life-history parameters has more desirable statistical properties. In our simulations, we found that the mRV estimates are slightly biased, but the bias is too small to be visible in the figures. mRV has the additional advantage that a sensitivity analysis (Caswell, 2019) can reveal how parameter uncertainty affects the precision of the estimate. In the field, such uncertainty can be considerably larger than in our simulations. Most importantly, we assumed that all stage transitions, including death, can be observed in the study population. In real populations, the estimation of, say, survival probabilities will often be unprecise and/or biased, as mortality cannot always be distinguished from emigration.

We purposely kept our model assumptions as simple as possible. We focussed on very simple life-histories, assuming that estimation issues that arise in simple scenarios will most likely be even worse in more complex life-histories. We assumed that our populations are closed (no emigration or immigration), that pedigree information is complete, and that the life-history parameters remained constant over time. Again, it is likely that RV estimates get worse when the situation is less ideal (as is the case for most field studies). Most of our simulations assume asexual reproduction, but as shown in Figure 5b they equally apply to populations with sexual reproduction and different sexes. However, they are not necessarily representative for more complicated situations, such as kin interactions, inbreeding or inbreeding avoidance, and sexual selection. But again, we would argue that methods that do not work well in a simple context will most likely also fail in more intricate situations.

Life history theory is one of the most advanced branches of evolutionary biology, with sophisticated tools and methods and a well-established fitness concept (Brommer, 2000). Individual-based simulations are rarely used in life-history studies, perhaps because analytical techniques are readily available. Yet, such simulations can provide valuable additional information. First, simulation models are very flexible and can easily be tailored to the intricacies encountered in real-world situations. For example, it is quite difficult to include sex- and age-structure, non-random mating, and kin interactions in a life history model without compromising analytical tractability, while this is straightforward in a simulation approach. Second, running replicated simulations gives researchers a good idea on the kind and degree of variation to be expected. Individual-based simulations are therefore a useful tool for judging the validity of empirical methods, such as the pedigree-based estimation of reproductive values.

## Acknowledgements

We thank the MARM research group, the Seychelles warbler research group and the reviewers of a previous version of this manuscript for their insightful comments on our study and the manuscript.

## Funding

MJB was funded by ALW NWO Grant No. ALWOP.531 awarded to JK and DR. The work of FJW is supported by the European Research council (ERC Advanced Grant No. 789240). The authors had no conflict of interest.

## Data availability

All data is archived on the University of Groningen Dataverse, which will be publicly accessible once this manuscript is published.

## Notes

### Competing Interest Statement

The authors have declared no competing interest.

### Summary of Updates

Simulations were altered so that both the mRV and pRV were estimated within the same simulation. We also removed the part on fluctuating environments and added a section on sexual reproduction and differences between sexes.

