## Supplementary materials 2022-11-03 for "Putting life history theory to the test - the estimation of reproductive values from field data"

### Supplementary Information

This supplement contains three appendices:

Appendix A:  Calculation of expected pRV

Appendix B:  The calculation of "true" RVs in Scenarios 1 to 4

Appendix C:  Effects of population size and time horizon on the pedigree-based estimation of RV

### Appendix A: Calculation of expected pRV

In this appendix, we derive an expression for $pv_j(t)$, the expected *per capita* number of descendants of individuals that were in stage *j* at time $t=0$. For a pedigree of depth *t*, the realized value of $pv_j(t)$ estimates the "pedigree reproductive value (pRV)" of stage *j*. By definition of "reproductive value", $pv_j(t)$ is expected to converge to the RV of stage *j* for large values of *t*.

Consider a life history model that is characterized by a matrix **A** (see eqn (1) in the main text). By definition, matrix element $a_{ij}$ is the *per capita* contribution of a member of stage *j* to the number of individuals in stage *i* in the next time step. Let us define $pv_{ij}(t)$ as the expected number of descendants that are in stage *i* at time *t* of an individual that was in stage *j* at time $t=0$. Obviously,

$$pv_j(t)=\sum_i pv_{ij}(t)\,. \qquad \text{(A1)}$$

By definition of $pv_{ij}(t)$ and $a_{ij}$, $pv_{ij}(1)$ and $pv_j(1)$ are given by

$$pv_{ij}(1)=a_{ij} \text{ and } pv_j(1)=\sum_i a_{ij}\,. \qquad \text{(A2)}$$

In other words, $pv_j(1)$ is just the sum of the elements in the *j*th column of the life history matrix **A**. It is now straightforward to derive a recurrence equation for the terms $pv_{ij}(t)$:

$$pv_{ij}(t+1)=\sum_k a_{ik}\cdot pv_{kj}(t)\,. \qquad \text{(A3)}$$

In words: for an individual that was in stage *i* at time $t=0$, the expected number of descendants in stage *i* at time $t+1$ is given by the sum of descendants in stage *k* at time *t* [$pv_{kj}(t)$] times the *per capita* contribution of these descendants to stage *i* [$a_{ik}$] in the following time step. Mathematically speaking, $pv_{ij}(t+1)$ is given by the scalar product (or "dot product") of the vector $\mathbf{pv}_{\cdot j}(t)$ and the *i*th row of matrix **A**. Equation (A3) specifies a linear system of recurrence equations with constant coefficients. In principle, such a system is analytically solvable, but for our purposes the solution does not inspire much insight. It is easier to iterate system (A3), with starting condition (A2). This yields all values $pv_{ij}(t)$ and, via (A1) the values of $pv_j(t)$ that we wanted to derive.

Figure A1 and A2 show that the iteratively calculated values of $pv_j(t)$ agree very well with the median pRV of 100 replicate simulations at time *t*. We can conclude that for relatively small values of *t* ($t$ < 12 in Fig. A1(a), $t$ < 5 in Fig. A1(b), $t<20$ in Fig. A2(a) and $t\leq 5$ in Fig. A2(b)) the *expected* values of the pRVs differ systematically from the "true" RVs. For large values of *t*, the expected values of the pRVs converge to the true RVs. However, as illustrated in Figures 3, 4 and 5, the pRVs from individual simulations (corresponding to a single study population) can differ enormously from the expected values, even if the pedigree is based on a large number of individuals and the pedigree is very deep (see Appendix C).

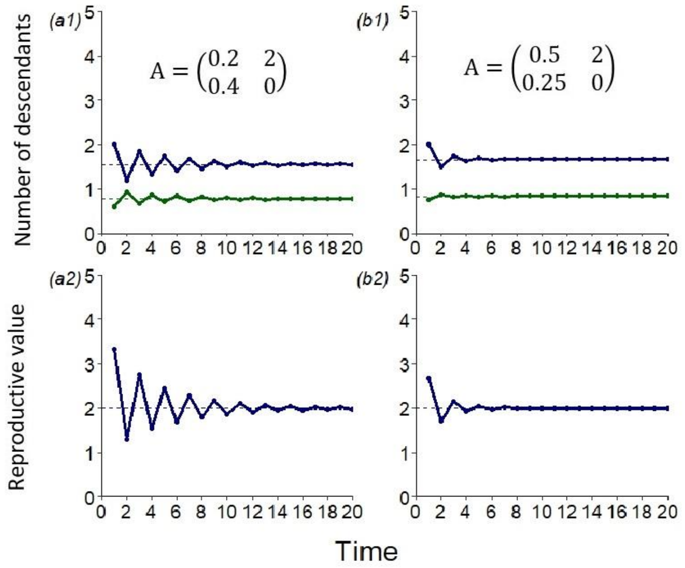


**Figure A1. Expected pRV after *t* time steps in Scenario 1.** For two parameter combinations of the life-history model with two age classes (see inset) panels (a1) and (b1) show the expected number of descendants after *t* time steps [$pv_j(t)$], as iteratively calculated on the basis of eqns (A1) to (A3). Panels (a2) and (b2) show the (normalised) pRV values, which are obtained by scaling $pv_1(t)$ to one. Notice that convergence is much slower for the parameters in (a) than for the parameters in (b). A comparison of panel (b2) with Figure 4e reveals a close agreement of the expected values with the median of 100 replicate simulations.

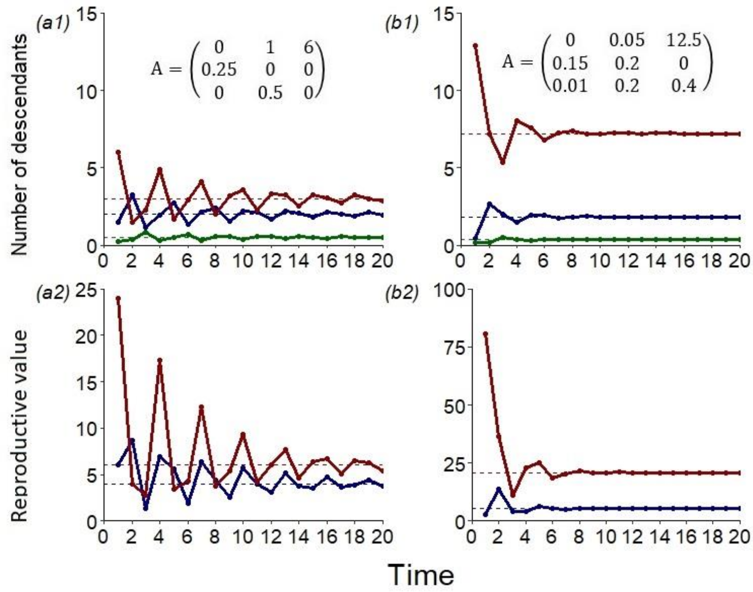


**Figure A2. Expected pRV after *t* time steps in Scenario 2.** For two parameter combinations of the life-history model with three age classes (see inset) panels (a1) and (b1) show the expected number of descendants after *t* time steps [$pv_j(t)$], as iteratively calculated on the basis of eqns (A1) to (A3). Panels (a2) and (b2) show the (normalised) pRV values, which are obtained by scaling $v_1$ to one. A comparison of panel (a1) with Figure 2c and of panel (b2) with Figure 5a reveals a close agreement of the expected values with the median pRV of 100 simulations.

### Appendix B: The calculation of "true" RVs in Scenarios 1 to 4

Scenario 1: A population with two age classes.

This scenario is characterized by the life-cycle graph in Fig. 1a or, equivalently, by the Leslie matrix:

$$\mathbf{A} = \begin{pmatrix} F_1 & F_2 \\ P_1 & 0 \end{pmatrix}. \qquad \text{(B1)}$$

At demographic and ecological (λ=1) equilibrium, the reproductive values are given by the vector equation $\mathbf{v}^T = \mathbf{v}^T \cdot \mathbf{A}$, which corresponds to a system of two equations:

$$\begin{aligned} v_1 &= F_1 \cdot v_1 + P_1 \cdot v_2 \\ v_2 &= F_2 \cdot v_1 \end{aligned}. \qquad \text{(B2)}$$

Normalizing reproductive values by setting $v_1 = 1$, this system yields eqns (7a) and (7b) in the text:

$$\begin{aligned} v_2 &= F_2 \\ F_1 + P_1 \cdot F_2 &= 1 \end{aligned}. \qquad \text{(B3)}$$

These equations make perfect sense: the reproductive value of the final age class is always equal to the expected number of offspring produced at this age class (see also Scenario 2); and the left-hand side of the second equation corresponds to expected lifetime reproductive success, which needs to be equal to 1 at ecological equilibrium.

Scenario 2: A population with three age classes.

This scenario is characterized by the life-cycle graph in Fig. 1b or, equivalently, by the Leslie matrix:

$$\mathbf{A} = \begin{pmatrix} 0 & F_2 & F_3 \\ P_1 & 0 & 0 \\ 0 & P_2 & 0 \end{pmatrix}. \qquad \text{(B4)}$$

At demographic and ecological (λ=1) equilibrium, the reproductive values are given by:

$$\begin{aligned} v_1 &= P_1 \cdot v_2 \\ v_2 &= F_2 \cdot v_1 + P_2 \cdot v_3 \\ v_3 &= F_3 \cdot v_1 \end{aligned}. \qquad \text{(B5)}$$

After normalizing ($v_1 = 1$), the solution of this system yields eqns (8a) and (8b) in the main text:

$$\begin{aligned} &v_2 = F_2 + P_2 F_3,\ v_3 = F_3 \\ &P_1 \cdot (F_2 + P_2 F_2) = 1 \end{aligned}. \qquad \text{(B6)}$$

As in Scenario 1, the left-hand side of the second equation corresponds to expected lifetime reproductive success, which needs to be equal to 1 at ecological equilibrium.

Scenario 3: A population with three behavioural classes.

This scenario is characterized by the life-cycle graph in Fig. 1c or, equivalently, by the stage transition matrix:

$$\mathbf{A} = \begin{pmatrix} P_j & F_h & F_b \\ T_{hj} & P_h & T_{hb} \\ T_{bj} & T_{bh} & P_b \end{pmatrix}. \quad \text{(B7)}$$

Now, the reproductive values at demographic and ecological (λ=1) equilibrium are given by:

$$\begin{aligned} v_j &= P_j \cdot v_j + T_{hj} \cdot v_h + T_{bj} \cdot v_b \\ v_h &= F_h \cdot v_j + P_h \cdot v_h + T_{bh} \cdot v_b \\ v_b &= F_b \cdot v_j + T_{hb} \cdot v_h + P_b \cdot v_b \end{aligned} . \quad \text{(B8)}$$

After normalizing ( $v_j = 1$ ), this system can readily be solved, yielding explicit equations for $v_h$ and $v_b$ and a consistency requirement for ecological stability (a stationary population). As these equations do not inspire much insight, they are not reproduced here.

Scenario 4: A population with sex and age classes.

This scenario, which is characterized by the life-cycle graph in Fig. 1d, is more complicated. If we order the states as follows: young females, young males, adult females, adult males, the stage transition matrix is of the form:

$$\mathbf{A} = \begin{pmatrix} \frac{1}{2}(1-s)F_{fy} & \frac{1}{2}(1-s)F_m & \frac{1}{2}(1-s)F_{fa} & \frac{1}{2}(1-s)F_m \\ \frac{1}{2}sF_{fy} & \frac{1}{2}sF_m & \frac{1}{2}sF_{fa} & \frac{1}{2}sF_m \\ P_{fy} & 0 & P_{fa} & 0 \\ 0 & P_{my} & 0 & P_{ma} \end{pmatrix}. \quad \text{(B9)}$$

The last two rows correspond to survival and the transition from young to adult stage. For example, the matrix element $a_{13} = P_{fy}$ indicates the probability that a young female survives, which implies that she moves to the class of adult females. The first two rows correspond to the production of offspring in the four life-history states. The factor $\frac{1}{2}$ indicates that, in a diploid population, per parent only half of the genes are transmitted to the offspring. Alternatively, one might say that half of each newly produced offspring is ascribed to the mother and half to the father (otherwise, each offspring would be counted twice. The factors 1-*s* and *s* indicate that a newly produced offspring is a young female with probability 1-*s* and a young male with probability *s*. Hence, matrix elements $a_{11} = \frac{1}{2}(1-s)F_{fy}$ and $a_{12} = \frac{1}{2}sF_{fy}$ indicate that a young female contributes on average $\frac{1}{2}(1-s)F_{fy}$ young females and $\frac{1}{2}sF_{fy}$ young males to the next time step. We still need to define the term $F_m$, the average number of offspring produced per male (remember that we assumed that young and adult males do not differ in their probability to fertilize a female, and, hence, in their reproductive output). $F_m$ can be calculated by making use of the "Fisher condition" (Fisher, 1930), which states that the total reproductive output of all males must exactly match the total reproductive output of all females (because each offspring has one father and one mother). If $n_{fy}, n_{my}, n_{fa}$, and $n_{ma}$ denote the number of young females, young males, adult females, and adult males at demographic equilibrium, the Fisher condition is:

$$(n_{my} + n_{ma}) \cdot F_m = n_{fy} \cdot F_{fy} + n_{fa} \cdot F_{fa}, \tag{B10}$$

yielding:

$$F_m = \frac{n_{fy} \cdot F_{fy} + n_{fa} \cdot F_{fa}}{n_{my} + n_{ma}}. \tag{B11}$$

The vector $(n_{fy}, n_{my}, n_f, n_{ma})$ corresponds to the stable stage distribution of the model, which is given by the right eigenvector of the matrix **A** in (B9). It is, however, easier to derive the stable stage distribution from the corresponding demographic model where all reproduction is ascribed to the females. This model is characterized by the following matrix:

$$\mathbf{B} = \begin{pmatrix} (1-s)F_{fy} & 0 & (1-s)F_{fa} & 0 \\ sF_{fy} & 0 & sF_{fa} & 0 \\ P_{fy} & 0 & P_{fa} & 0 \\ 0 & P_{my} & 0 & P_{ma} \end{pmatrix}. \tag{B12}$$

The stable stage distribution of **B** (which is identical to the stable stage distribution of **A**) is given by the eigenvector equation $\mathbf{n} = \mathbf{B} \cdot \mathbf{n}$, which is equivalent to the system of equations:

$$\begin{aligned} n_{fy} &= (1-s) \cdot (F_{fy}\, n_{fy} + F_{fa}\, n_{fa}) \\ n_{my} &= s \cdot (F_{fy}\, n_{fy} + F_{fa}\, n_{fa}) \\ n_{fa} &= P_{fy}\, n_{fy} + P_{fa}\, n_{fa} \\ n_{ma} &= P_{my}\, n_{my} + P_{ma}\, n_{ma} \end{aligned}. \tag{B13}$$

The 2$^{nd}$ equation implies $F_{fy}\, n_{fy} + F_{fa}\, n_{fa} = n_{my}/s$, while the 4$^{th}$ equation gives $n_{ma} = (P_{my}/(1-P_{ma})) \cdot n_{my}$. Inserting these expressions into (B11) and simplifying yields an expression of $F_m$ in terms of the model parameters:

$$F_m = \frac{1}{s} \cdot \frac{1-P_{ma}}{1-P_{ma}+P_{my}}. \tag{B14}$$

Now we know $F_m$, we can focus on the eigenvector equation $\mathbf{v}^T = \mathbf{v}^T \cdot \mathbf{A}$ yielding the reproductive values. In the present scenario, it corresponds to a system of four equations:

$$\begin{aligned} v_{fy} &= \tfrac{1}{2} F_{fy} \cdot \left((1-s) \cdot v_{fy} + s \cdot v_{my}\right) + P_{fy} \cdot v_{fa} \\ v_{my} &= \tfrac{1}{2} F_m \cdot \left((1-s) \cdot v_{fy} + s \cdot v_{my}\right) + P_{my} \cdot v_{ma} \\ v_{fa} &= \tfrac{1}{2} F_{fa} \cdot \left((1-s) \cdot v_{fy} + s \cdot v_{my}\right) + P_{fa} \cdot v_{fa} \\ v_{ma} &= \tfrac{1}{2} F_m \cdot \left((1-s) \cdot v_{fy} + s \cdot v_{my}\right) + P_{ma} \cdot v_{ma} \end{aligned}. \tag{B15}$$

The joint term in these equations is the average reproductive value of juveniles, which we can normalize to one:

$$\bar{v}_j = (1-s)\cdot v_{fy} + s\cdot v_{my} = 1\,, \qquad \text{(B16)}$$

which simplifies (B15) considerably. The last two equations give us immediately the reproductive values of the two adult states:

$$v_{fa} = \tfrac{1}{2}\cdot\left(F_{fa}/(1-P_{fa})\right),\quad v_{ma} = \tfrac{1}{2}\cdot\left(F_m/(1-P_{ma})\right). \qquad \text{(B17)}$$

These equations make perfect sense, as the terms in brackets corresponds to the expected lifetime reproductive success of an adult female and an adult male, respectively. Inserting the expression for $v_{ma}$ into the second equation of (B15) yields (by making use of (B14)) a simple equation for $v_{my}$:

$$v_{my} = \tfrac{1}{2}F_m + P_{my}\cdot\tfrac{1}{2}\cdot\left(F_m/(1-P_{ma})\right) = \tfrac{1}{2}F_m\cdot\left(1+(P_{my}/(1-P_{ma}))\right) = \tfrac{1}{2}F_m\cdot\left(1/(sF_m)\right) = 1/(2s)\,.$$

Inserting this expression into (B16) gives us the reproductive values of the two juvenile states:

$$v_{fy} = \tfrac{1}{2}\cdot\tfrac{1}{1-s},\quad v_{my} = \tfrac{1}{2}\cdot\tfrac{1}{s}, \qquad \text{(B18)}$$

which again makes perfect sense, because the reproductive values of juvenile males and females are inversely proportional to the abundance of juvenile males and females, respectively.

Notice that $v_{fy}$ can also be determined differently, by inserting the expression for $v_{fa}$ in (B17) into the first equation of (B15):

$$v_{fy} = \tfrac{1}{2}F_{fy} + P_{fy}\cdot\tfrac{1}{2}\cdot\left(F_{fa}/(1-P_{fa})\right) = \tfrac{1}{2}\cdot\left(F_{fy} + F_{fa}\cdot(P_{fy}/(1-P_{fa}))\right).$$

Equating this expression for $v_{fy}$ with the one in (B18) yields the consistency requirement:

$$(1-s)\cdot\left(F_{fy} + F_{fa}\cdot(P_{fy}/(1-P_{fa}))\right) = 1\,. \qquad \text{(B19)}$$

The population is in ecological equilibrium (λ=1) if (and only if) condition (B19) is satisfied.

Please note that for consistency throughout the manuscript, we used a different normalisation in Figure 5b: instead of setting the average reproductive value of juveniles to 1 (as in (B16)), we set the reproductive value of the first class (young females) to 1.

### Appendix C: Effects of population size and time horizon on pRV

In the main text, we focused on a time horizon of 20 time steps, as the corresponding graphs show more clearly the systematic deviation of the pedigree-based estimates of the RVs from their true values (that was discussed in Appendix A). Moreover, all simulations shown in the main text are based on populations consisting of 1000 individuals. Here we illustrate that our conclusions on pRV in the main text also apply to a time horizon of 100 time steps and to different population sizes.

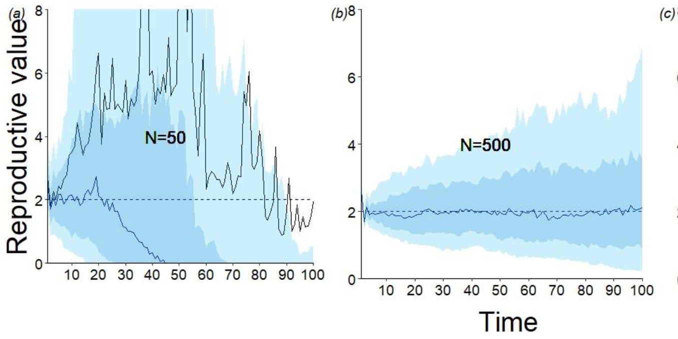


**Figure C1. Effect of population size and time horizon on pRV estimates in Scenario 1 (two age classes).** This figure is the counterpart to Fig. 4e in the main text. For the same life-history parameters ($F_1 = 0.5$, $P_1 = 0.25$, $F_2 = 2$), 100 replicate simulations were run for different population sizes and a longer time horizon. (a) In case of a small population ($N = 50$), the gene-dropping method fails on a longer time scale due to demographic stochasticity: after some time, all descendants can either be traced back to age-one or to age-two individuals in the original population, while there are no descendants left from the other category. (b,c) For $N = 500$ and $N = 5{,}000$ the variation across simulations increases with the time horizon.

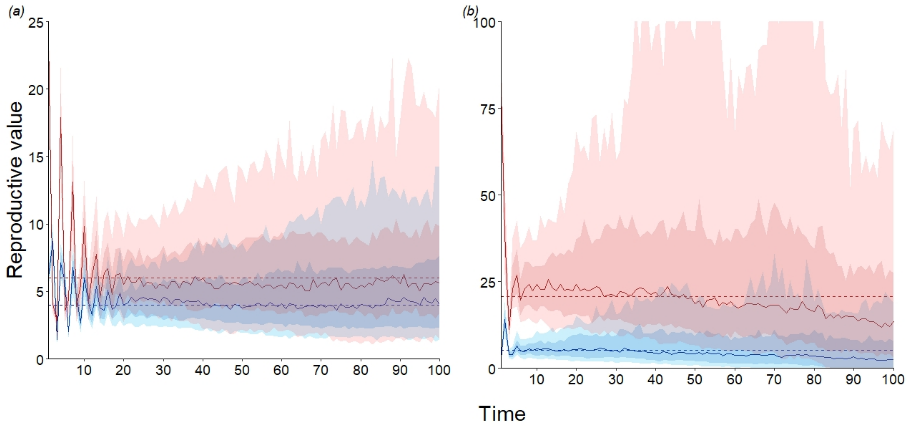


**Figure C2. Effect of time horizon on pRV estimates in (a) Scenario 2 (3 age classes) and (b) Scenario 3 (3 behavioural classes).** Extensions of the simulations on Scenario 2 (Fig. 3e, parameters as in Fig. B2a) and Scenario 3 (Fig. 5a, same parameters) from 20 to 100 time steps. Again, the variation across simulations increases with the time horizon. Notice that eventually the median of the 100 pRV values does no longer agree with the true RV, due to demographic stochasticity.
